# A new yeast-based bioreporter for simple, sensitive, and cost-effective detection of dioxin-like compounds

**DOI:** 10.1101/2024.06.30.601204

**Authors:** Yingying Liang, Hailin Liu, Lin Wang, Jing Zhao, Shunyi Li, Li Yi, Sijing Jiang, Zhenghui Lu, Guimin Zhang

## Abstract

Dioxin-like compounds (DLCs) are environmental xenobiotics that can activate the aryl hydrocarbon receptors (AhR), thereby imposing a significant threat to human health through biomagnifications processes. In this study, a dioxin-activated nano-luminescent *Saccharomyces cerevisiae* bioreporter, called DnaSc, was developed for simple and rapid detection of DLCs and AhR agonists. The bioreporter used nano-luciferase NLuc as a signal generator to emit bioluminescent signals in response to DLCs without cell lysis. Through optimizing ARNT expression and engineering the aryl hydrocarbon receptor (AhR), the yeast-based bioassay exhibited a detection limit of 10 fM for 2,3,7,8-tetrachlorodibenzo-p-dioxin (TCDD) within 6 h, making it the most sensitive whole-cell biosensor reported to date. Furthermore, the detection capacity of the DnaSc bioassay for DLCs and AhR agonists was characterized. In summary, the yeast-based bioreporter developed in this study provided a simple, sensitive, and cost-effective method for DLCs detection.

## 1 Introduction

Polychlorinated dibenzo-p-dioxins (PCDDs) and polychlorinated dibenzofurans (PCDFs), collectively known as “dioxins”, are a type of compounds with similar chemical structures and physicochemical properties [1, 2]. There are 75 PCDDs and 135 PCDFs, and the main difference between them is the position and number of chlorine atoms [3]. TCDD in the PCDDs group is considered as one of the most toxic pollutants known to date [4, 5]. Dioxins are principally released into the atmosphere during processes such as municipal and medical solid waste incineration, chlorine bleaching of paper and wood pulp, chemical metallurgy, and cement production [6-8]. Due to their persistence in the environment and bioaccumulation in the food chain, dioxins accumulate easily in human tissues through the processes of biomagnification, posing a significant threat to human health [4, 9].

The routine method for quantifying dioxin concentrations is the high-resolution gas-chromatography coupled to high-resolution mass-spectrometry (HRGC-HRMS), which is widely accepted in official international standard methods [10-12]. Although highly sensitive and selective, HRGC-HRMS suffers from high operational costs because of the requirement of sophisticated instruments, well-trained operators, and expensive chemicals [13, 14].

The cytosolic aryl hydrocarbon receptor (AhR) acts as the receptor of dioxin-like compounds (DLCs) [15, 16]. After binding to DLCs, AhR translocates to the nucleus, where it dimerizes with the AhR nuclear translocator (ARNT) to recognize and bind to consensus regulatory DNA sequences called xenobiotic-responsive elements (XREs) in the enhancer regions of responsive genes and activates their expression [17, 18]. Based on this mechanism, various types of bioassays have been developed as less expensive and complex alternatives for DLC screening. The chemically activated luciferase expression (CALUX) is the most widely adopted bioassay for DLCs detection, which uses a transgenic cell line (U.S. Patent No. 5854010) to express luciferase in response to the binding of DLCs to AhR. CALUX has been approved by the United States Environmental Protection Agency (EPA) and is widely adopted in the United States (FDA and EPA), the European Union, Chile, Japan, and other countries as a new rapid detection technology for screening dioxin-like pollutants in various environmental samples, including soil, water, flue gas, sludge, feed, additives, and biological tissues [19, 20].

*Saccharomyces cerevisiae* has also been used as a host to develop bioassays for the detection of DLCs due to its rapid growth, cost-effectiveness, and stability. Miller developed the CROMIS AhR kit using *S. cerevisiae* W303 as a host and β-galactosidase lacZ as a reporter [21]. Xu et al created a DLC-responsive autobioluminescent *S. cerevisiae* bioreporter BLYAhS through functionally expressing a synthetic bacterial luciferase reporter gene cassette (*luxCDABEfrp*), which can continuously produce bioluminescent signal without the need for cell lysis. The limit of detection (LOD) of BLYAhS for the model dioxin 2,3,7,8-tetrachlorodibenzo-p-dioxin (TCDD) is 500 pM [22].

Nano-luciferase (NLuc) is a newly designed luciferase with advantages such as strong luminescence, small molecular weight, no requirement for ATP, and stable physicochemical properties at various pH values and temperatures. These properties make NLuc an attractive reporter gene for sensitive bioluminescence detection [23]. In this study, a dioxin-activated nano-luminescent *S. cerevisiae* bioreporter DnaSc was constructed for the detection of DLCs using NLuc as a signal generator without the need for cell lysis. In addition, by optimizing the AhR and ARNT elements in the sensing module, the bioreporter achieved a remarkable detection limit of 10 fM for TCDD within 6 hours.

## 2 Materials and Methods

### 2.1 Enzymes and reagents

Restriction endonucleases were purchased from Takara (China) and NEB (USA). DNA polymerases, plasmid mini-prep kit (Cat. DC201-01), PCR purification kit (Cat. DC301-01), and DNA cloning kit (Cat. C112-01) were all obtained from Vazyme Biotech Co., Ltd (China). Nano-Glo Live Cell Assay System kit was purchased from Promega (Beijing) Biotech Co., Ltd; yeast nitrogen base (YNB) and DO Supplement-Trp were purchased from Beijing Coolaber Technology Co., Ltd (China). Tryptone and yeast extract were obtained from OXOID (United Kingdom) and Casamino acids were purchased from BD Corporation (USA). TCDD was purchased from Wellington Laboratories Inc (Canada). β-NF, and 2-CBP was obtained from Aladdin (China). Other routine reagents were purchased from China National Pharmaceutical Group Chemical Reagent Co., Ltd.

### 2.2 Plasmid Construction

To construct the sensing module, human AhR and ARNT cDNA fragments were cloned into the *Not*I and *Bam*HI restriction sites of pESD, respectively [24] to obtain pESD-AHRC (pESD-AhR-P_GAL1, 10_-ARNT). To construct the reporter module, five dioxin-responsive elements XRE (XRE5) were fused to the promoter PCYC1 by overlapping PCR to generate the dioxin-responsive promoter XRE5-PCYC1. XRE5-PCYC1 and the reporter gene NLuc were then cloned into pESD to produce the reporter plasmid pESD-XRE5-PCYC1-Nluc. For ARNT overexpression, the P_GAL10_-ARNT expression cassette was integrated into *Bgl*II and the reporter plasmid pESD-XRE5-PCYC1-NLuc was double digested with *Sac*I to obtain the ARNT overexpression and reporter plasmid pESD-XRE5-Nluc-ARNT. To construct the AhR mutant V381A, whole plasmid PCR was conducted using plasmid pESD-AHR-ARNT-G as a template and primer pair AHR381-F/AHR381-R (listed in Table S1). The PCR products were digested with *Dpn*I and transformed into *E. coli* DH5α to produce plasmid pESD-AHRC381. All plasmid constructions were carried out using *Escherichia coli* strain DH5α.

### 2.3 Yeast Strain Construction

The sensing module (AhR-P_GAL1,10_-ARNT) was integrated into the intergenic region between the DTP1 and the YCL49C genes of the yeast strain EBY100 [24]; a DNA fragment containing AhR-P_GAL1,10_-ARNT, the G418 resistance gene, and approximately 200 bp of H-up and H-down homologous arms was generated by overlapping PCR. Two *Xho*I restriction sites were introduced at the N-terminus of H-up and the C-terminus of H-down, respectively. The purified DNA fragment H-up-ARNT-P_GAL1,10_-AhR-G418-H-down was cloned into the pESD vector double-digested with *EcoR*I and *BamH*I to obtain the plasmid pESD-AHR-ARNT-G, which was further digested with *Xho*I to obtain a linearized fragment and electroporated into yeast EBY100 (Fig. S1). The transformants were cultivated on YPD plates containing geneticin G418, and further verified by colony PCR and DNA sequencing to obtain the recombinant yeast strain EBY100-AHRC. Then, the reporter plasmid pESD-XRE5-Nluc was transformed into EBY100-AHRC via electroporation. The resultant strain (AHRC) was used for DLCs bioassay. To detect the effect of ARNT overexpression on the detection performance of DLCs, pESD-XRE5-Nluc-ARNT was transformed into EBY100-AHRC to obtain strain AHRC-ARNT. To confirm the effect of AHR381 on the detection performance of DLCs, plasmid pESD-AHRC381 was transformed into EBY100 to obtain EBY100-AHRC381, and then the reporter plasmids pESD-XRE5-Nluc and pESD-XRE5-Nluc-ARNT were transformed into EBY100-AHRC381 to get strain AHRC381 and DnaSc, respectively.

### 2.4 TCDD Sensing Assay

Yeast bioassay strains were cultivated in YNB-CAA-Glu medium (0.67 g YNB, 2 g Glucose,0.5 g Casamino acids for 100 mL) with shaking at 30 °C for approximately 12 h. Cells were collected by centrifugation and washed twice with sterile water. The cell pellets were resuspended in YNB-CAA-Gal liquid media (0.67 g YNB,2 g Galactose,0.5 g Casamino acids for 100 mL) containing different concentrations of TCDD until the OD600 cell density reached 0.3. After 6 hours of cultivation at 30 °C, 100 µL of the induced cell culture was mixed with 25 µL of NLuc substrate (Promega) consisting of Nano-Glo^®^ Live Cell Substrate and Nano-Glo^®^ LCS Dilution Buffer at a ratio of 1:19. Luminescent signals were immediately measured using a Glo Max 20/20 Luminometer (Promega).

## 3. Results

### 3.1 Construction and characterization of a yeast-based DLC bioassay system using NLuc as a reporter gene

To develop the sensing module, cDNA of human AhR and ARNT were co-expressed under the control of the GAL1-GAL10 bidirectional promoter (**Fig. 1A**). Subsequently, the sensing module was integrated into the chromosome of *S. cerevisiae* EBY100. As for the reporter module, the NLuc reporter gene was driven by the dioxin-responsive promoter XRE5-P_CYC1_, which contains five XRE sequence (5′-TCTTGCGTGACAAT-3′) located upstream of the P_CYC1_ minimal promoter. The expression cassette was cloned into the yeast expression plasmid pESD to obtain pESD-XRE5-P_CYC1_-NLuc (**Fig. 1A**), which was then transformed into *S. cerevisiae* EBY100 containing the sensing module to obtain the yeast-based DLC bioassay system (abbreviated as AHRC).

**Figure 1.**
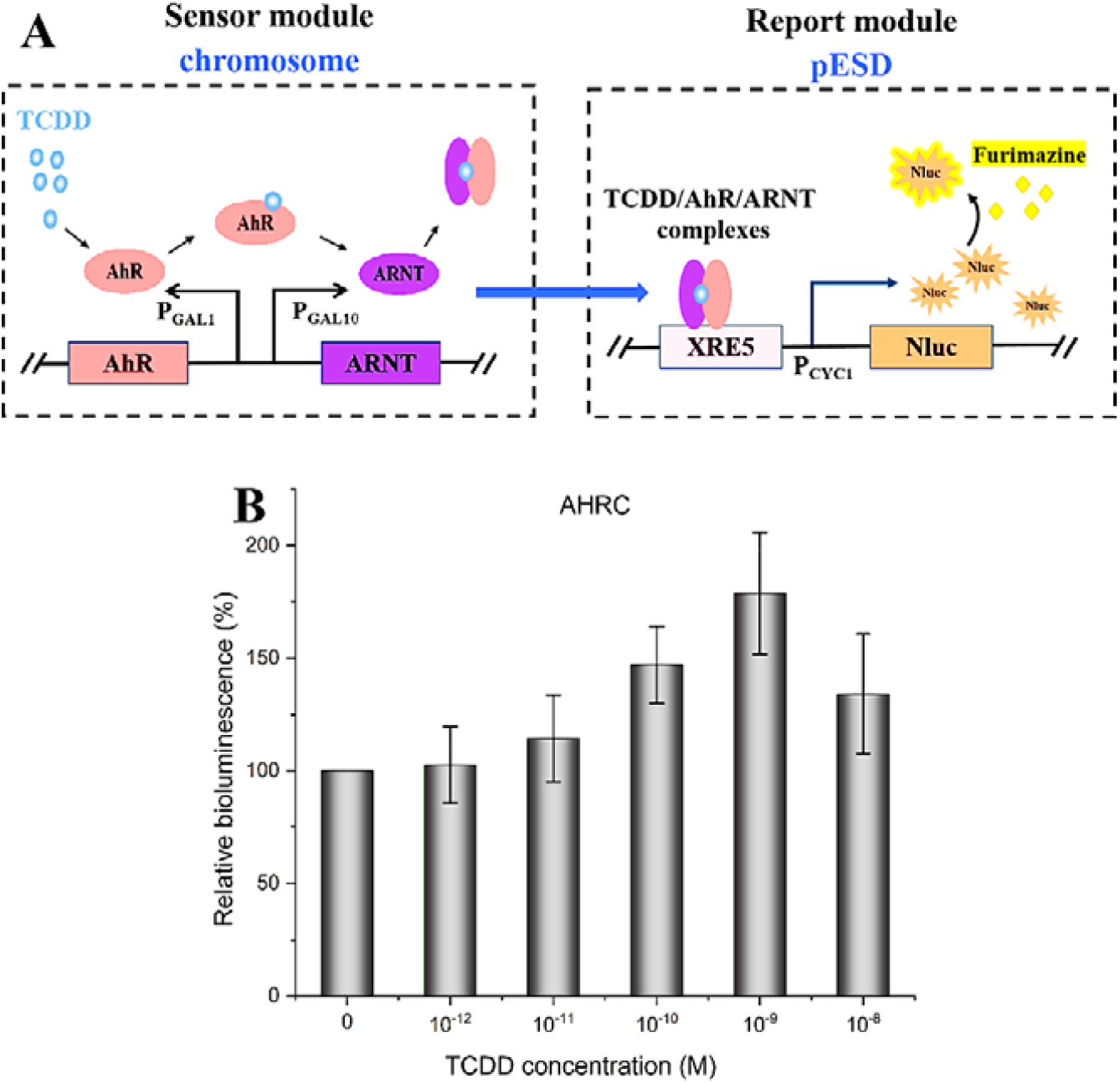
Yeast-based TCDD bioassay AHRC. A, Schematic diagram of the yeast-based DLC bioassay AHRC. The AHRC bioreporter strain was constructed by introducing the pESD-XRE5-P_CYC1_-NLuc reporter plasmid into the *S. cerevisiae* EBY100 strain harboring chromosomally integrated human AhR and ARNT genes under the control of the GAL1-GAL10 bidirectional promoter. B, Bioluminescent responses of *S. cerevisiae* AHRC reporter cells exposed to TCDD at concentrations ranging from 1 pM to 10 nM after 6 h induction.

The bioassay performance of AHRC was evaluated by exposing it to the model dioxin congener TCDD at concentrations ranging from 1 pM to 10 nM. As shown in **Figure 1B**, maximal induction was observed at 1 nM (10^−9^ M) TCDD, with a relative bioluminescent signal output of 170 % compared to the control. The LOD of AHRC for TCDD was determined to be 100 pM (10^−11^ M) after 6 h incubation.

### 3.2 Effects of ARNT overexpression on AHRC signaling outputs

In the presence of DLCs, AhR translocates to the nucleus and interacts with ARNT. The AhR/ARNT heterodimer then binds to XREs and activates the expression of downstream genes. Previous studies have shown that upregulation of ARNT expression increases the magnitude of the transcriptional response to TCDD in mice [25]. To understand the effect of ARNT expression on the performance of the yeast bioreporter AHRC bioassay, the P_GAL10_-ARNT expression cassette was integrated into the plasmid pESD-XRE5-P_CYC1_-NLuc (referred to as AHRC-ARNT) (**Fig. 2A**). As shown in **Figure 2B**, ARNT overexpression exerted a positive effect on the signal outputs, resulting in an approximately 30 % improvement in relative bioluminescence at 100 pM TCDD. Consequently, the detection limit of AHRC-ARNT for TCDD was reduced from 100 pM to 1 pM (10^−12^ M) (**Fig. 2B**).

**Figure 2.**
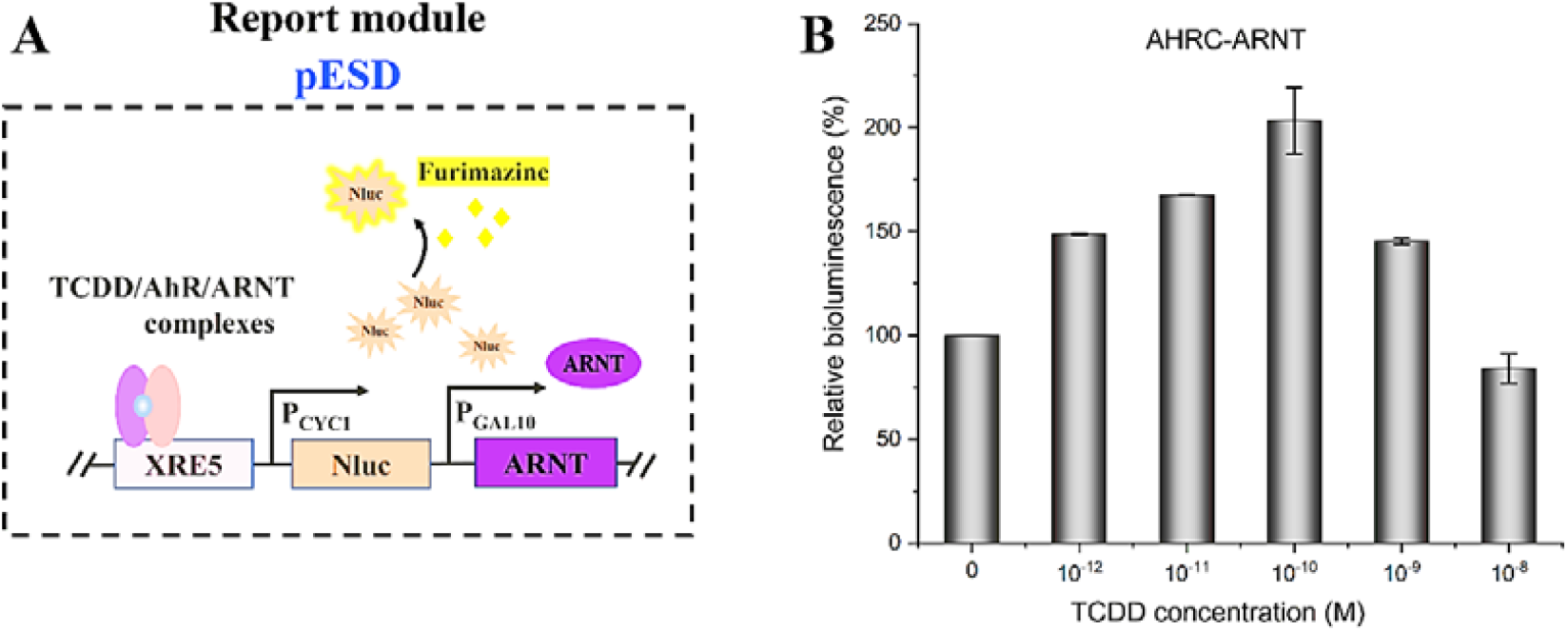
Yeast-based TCDD bioassay AHRC-ARNT. A, Schematic diagram of the yeast-based DLCs bioassay AHRC-ARNT. B, Bioluminescent responses of AHRC-ARNT reporter cells exposed to TCDD at concentrations ranging from 1 pM to 10 nM after 6 h induction.

### 3.3 Effect of the affinity between AhR and ligand on AHRC sensitivity

The affinity between AhR and exogenous ligands has been reported to be related to residue 381 in the ligand-binding pocket. Therefore, the valine residue at position 381 of the AhR protein in AHRC was substituted by the residue alanine (**Fig. 3A**). The resulting bioreporter, called the AHRC381, exhibited nearly the same maximal signal as AHRC. However, maximal induction occurred at 10 pM (10^−13^ M) TCDD, 100 times lower than AHRC (**Fig. 3B**). Similarity, the detection limit of TCDD by AHRC381 was reduced by 100-fold, from 100 pM to 1 pM (**Fig. 3B**).

**Figure 3.**
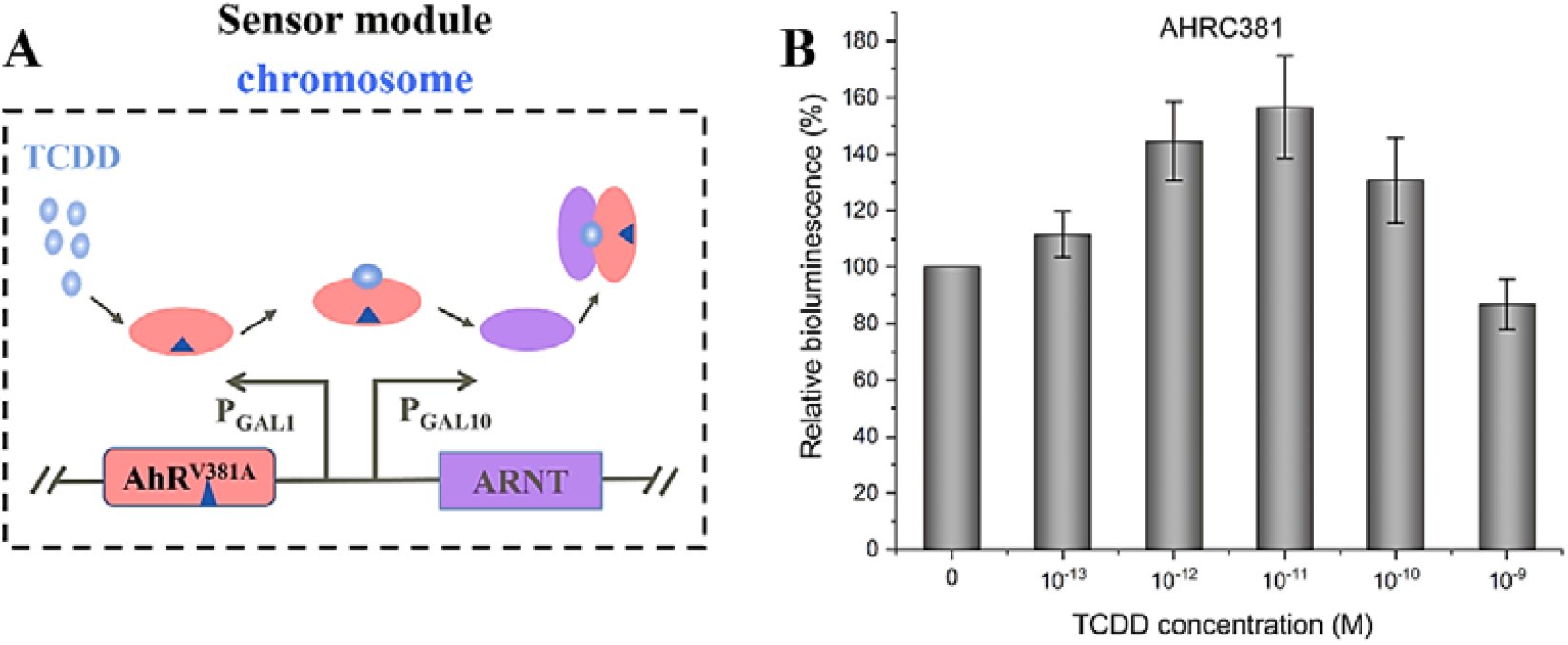
Yeast-based TCDD bioassay AHRC381. A, Schematic diagram of the yeast-based DLC bioassay AHRC381. B, Bioluminescent response of AHRC381 reporter cells exposed to TCDD at concentrations ranging from 0.1 pM to 1 nM after 6 h induction.

### 3.4 Generation of highly sensitive yeast-based DLC bioassay

Since both ARNT overexpression and AhR (V381A) mutant have positive impacts on the response and sensitivity of AHRC, by combining the beneficial effects of ARNT overexpression and AhR(V381A) mutation, we developed a dioxin-activated nano-luminescent *S. cerevisiae* bioreporter named DnaSc (**Fig. 4A**). Interestingly, the detection limit of TCDD by DnaSc was found to be significantly reduced to 10 fM (**Fig. 4B**), with maximal induction observed at 100 fM (10^−13^ M) TCDD, resulting in a relative bioluminescent signal output of 150 % compared to the control. Notably, DnaSc readily induced a clear bioluminescent signal when exposed to TCDD at concentrations ranging from 10 fM to 1 nM (**Fig. 4B**).

**Figure 4.**
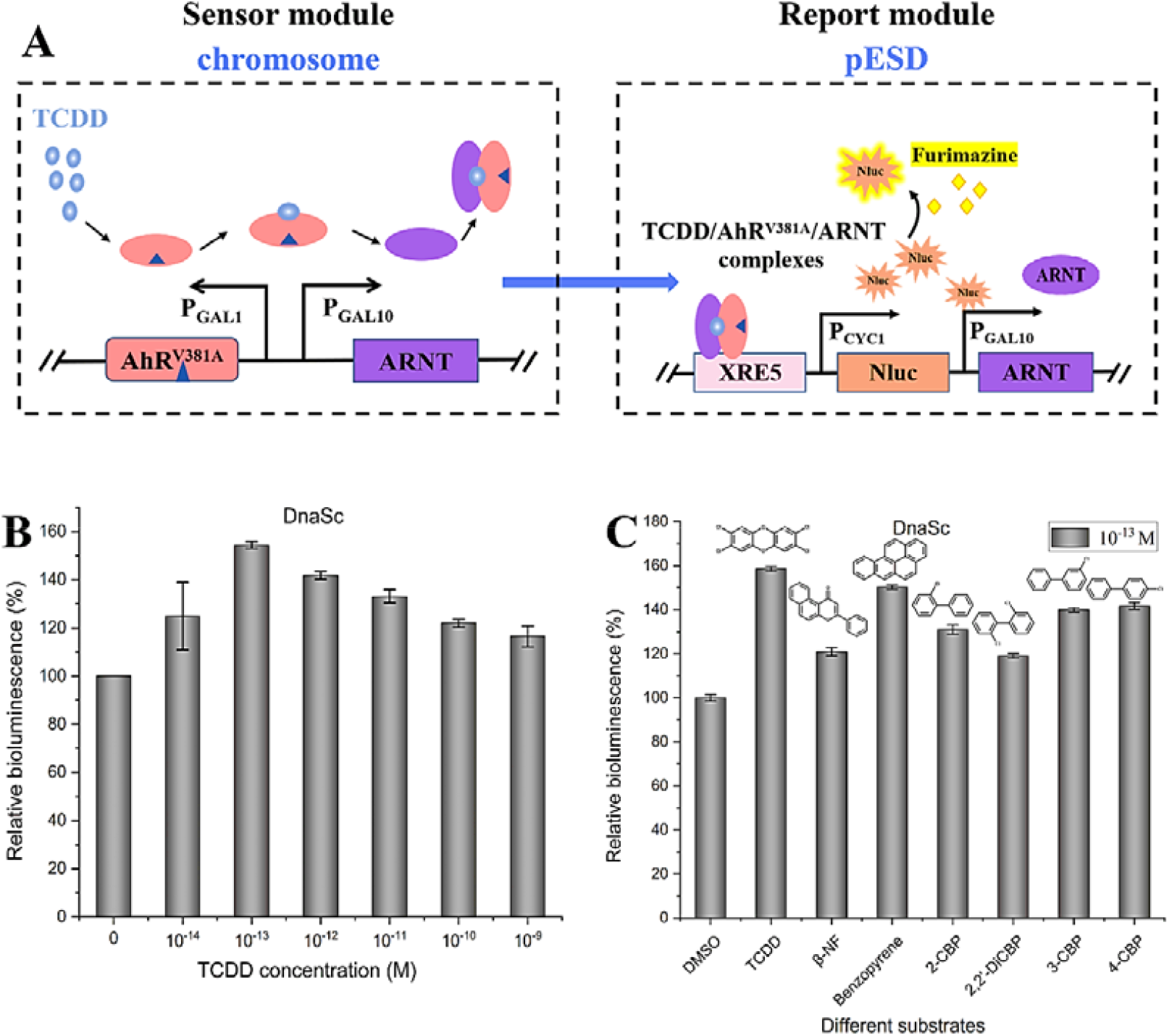
Yeast-based TCDD bioassay DnaSc. A, Schematic diagram of the yeast-based DLC bioassay DnaSc. B, Bioluminescent response of DnaSc reporter cells exposed to TCDD at concentrations ranging from 10 fM to 1 nM after 6 h induction. C. Detection of various DLCs and AhR agonists using the *S. cerevisiae* DnaSc bioreporter. Each DLC was tested using a concentration of 100 fM.

### 3.5 Detection of DLCs using DnaSc bioassay

DnaSc is able to detect a wide range of DLCs, including 2-chlorinated biphenyl (2-CBP), 2,2’-dichlorobiphenyl (2,2’-DiCBP), 3-chlorinated biphenyl (3-CBP), and 4-chlorinated biphenyl (4-CBP), and two known AhR agonists, benzopyrene (BaP) and β-naphthoflavone (β-NF), at a concentration of 0.1 pM. All DLCs tested induced a bioluminescent response within 6 h of exposure (**Fig. 4C**). Also, Benzopyrene, 3-CBP, and 4-CBP induced signal outputs comparable to that of TCDD, highlighting the versatility and sensitivity of DnaSc in the efficient detection of multiple dioxin-like compounds.

## 4 Discussion

Dioxin-like compounds (DLCs) are persistent organic pollutants known to cause a range of adverse metabolic, reproductive, immunologic, and neoplastic effects. Given the critical significance of DLCs screening, various analytical techniques, such as HRGC/HRMS and bioassays using DLC-responsive promoters to drive reporter gene expression have been developed [26]. Among the commonly used reporter genes in DLC screening bioassays, firefly luciferase and β-galactosidase are the most widely used in bioassays such as CALUX [27]. Mammalian cell-based CALUX bioassays typically exhibit detection limits ranging from 0.3-2 pM, while yeast-based bioassays using luciferase and β-galactosidase as reporters have detection limits of 930 pM and 300 pM, respectively [19, 21, 27]. To eliminate the need to lyse cells, Xu et al constructed a yeast-based auto-bioluminescent bioreporter capable of detecting TCDD with a detection limit of 500 pM [22]. In our study, we harnessed the power of nano-luciferase (NLuc) to create a new yeast-based DLCs bioassay, AHRC, which emitted bioluminescent signals in response to DLCs without the need for cell lysis. The detection limit of TCDD was observed to be 100 pM using AHRC.

AHR and ARNT are key components of the DLCs signaling pathway. Upregulation of ARNT expression positively the signal output and sensitivity of AHRC to TCDD, suggesting that the concentration of AhR in the AHRC reporter cells exceeded that of ARNT. As a result, the increased ARNT expression level could react with free AhR to form AhR-Arnt heterodimers, which then activated the NLuc reporter gene. The substitution of V381 of AhR in AHRC with alanine reduced the detection limit by 100-fold. Indeed, previous studies reported that AhR residue 381 differs between humans (V381) and Neandertals (A381), resulting in a higher EC50 activity of Neandertal AhR-induced Cyp1a1 activity than that observed for human AhR by 150-fold to 1000-fold [28]. Interestingly, the mouse AhR encodes A375, a residue homologous to A381 in Neandertal and other nonhuman primates. However, quantitative ligand binding studies revealed that the affinity of the A375 mouse AhR is only 10-fold higher than that of the human AhR [29]. These results suggested that, in addition to ligand binding, other factors such as nuclear translocation and ANRT affinity may also play a role in determining the transcriptional activity of AhR381 mutants.

Through upregulating ARNT expression and introducing mutations in the AhR ligand binding pocket, we successfully engineered the DLCs bioassay, DnaSc, exhibiting an unprecedented detection limit of 10 fM for TCDD. This highly improved sensitivity exceeded that of mammalian cell-based bioassays and highlighted a significant advance in low cost and high efficiency DLC assays. Although the individual impacts of ARNT overexpression and AhR engineering on DLCs response have been elucidated in mice, this combination is novel and their effects on DLCs sensing in yeast remain to be determined. The superior sensitivity of DnaSc revealed the critical role of the sensing module (AhR-ARNT signaling pathway) in DLCs bioassays. However, in the previously published DLC bioassays, the sensing module is often neglected, and the main focuses are on using different reporter genes to optimize the reporter module. Furthermore, the DLCs-induced bioluminescent response in DnaSc is dose-independent, consistent with other DLCs bioassays [30]. One possible explanation for this phenomenon is that at higher concentrations, DLCs may exert additional adverse effects on yeast cells.

## 5 Conclusion

After further optimization by AhR engineering and ARNT overexpression, we developed a new bioluminescent *S. cerevisiae* bioassay based on an NLuc reporter gene for screening DLCs. The resulting bioassay DnaSc exhibited a detection limit of 10 fM for TCDD and other AhR agonists. Since the NLuc substrate furimazine is cell permeable, cell lysing is not required before bioluminescent signal measurement. Therefore, this bioassay was easy to conduct, cost-effective, highly sensitive, and compatible with high-throughput screening such as 96-well plates. Taken together, the innovative advances presented in our study highlighted the great potential of the DnaSc bioreporter for detecting trace amounts of TCDD with excellent precision and efficacy, paving the way for routine environmental and human health monitoring.

## Acknowledgements

This work was supported by National Key R&D Program of China (No.2018YFA0901100), National Natural Science Foundation of China (32370101) and Fundamental Research Funds for the Central Universities (buctrc202131).

## Appendix A. Supplementary data

Supplementary data to this article can be found in the supporting information file

## Graphical abstract

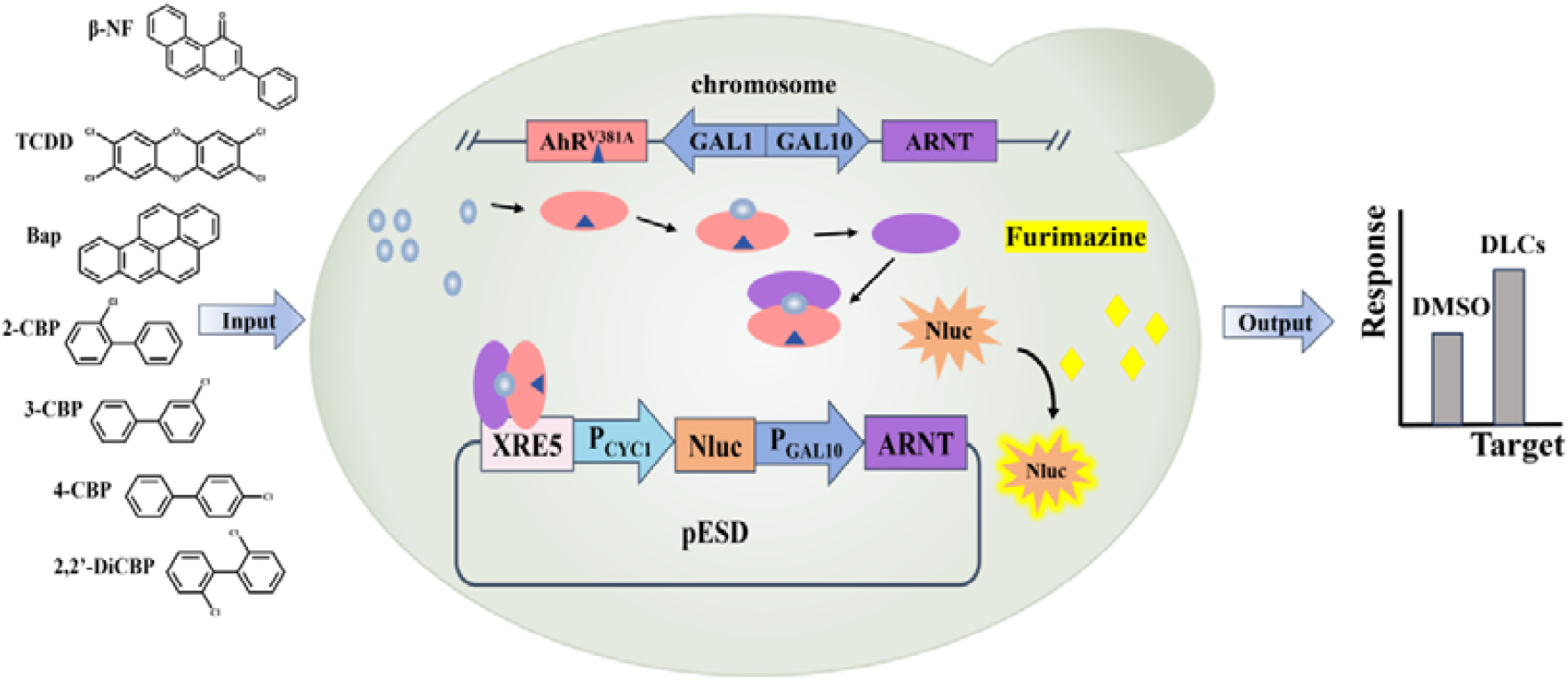

## References

[1] R. Fernández-González, I. Yebra-Pimentel, E. Martínez-Carballo, J. Simal-Gándara, A Critical Review about Human Exposure to Polychlorinated Dibenzo-p-Dioxins (PCDDs), Polychlorinated Dibenzofurans (PCDFs) and Polychlorinated Biphenyls (PCBs) through Foods, Crit Rev Food Sci Nutr 55(11) (2015) 1590–617.

[2] R. Lei, Z. Xu, Y. Xing, W. Liu, X. Wu, T. Jia, S. Sun, Y. He, Global status of dioxin emission and China’s role in reducing the emission, J Hazard Mater 418 (2021) 126265.

[3] B. Patrizi, M. Siciliani de Cumis, S. Viciani, F. D’Amato, Dioxin and Related Compound Detection: Perspectives for Optical Monitoring, Int J Mol Sci 20(11) (2019).

[4] E. Gaspard, P. Frenoy, D. Praud, T. Coudon, L. Grassot, A.A. Assi, B. Fervers, A. Gelot, F.R. Mancini, G. Severi, C. Besson, E. Faure, Association between cumulative airborne dioxin exposure and non-Hodgkin’s lymphoma risk in a nested case-control study within the French E3N cohort, Science of The Total Environment 906 (2024) 167330.

[5] E. Sørmo, K.M. Krahn, G.Ø. Flatabø, T. Hartnik, H.P.H. Arp, G. Cornelissen, Distribution of PAHs, PCBs, and PCDD/Fs in products from full-scale relevant pyrolysis of diverse contaminated organic waste, Journal of Hazardous Materials 461 (2024) 132546.

[6] S.V. Ajay, K.P. Prathish, Dioxins emissions from bio-medical waste incineration: A systematic review on emission factors, inventories, trends and health risk studies, J Hazard Mater 465 (2024) 133384.

[7] R. Lei, W. Liu, Y. He, T. Jia, C. Li, W. Su, Y. Xing, Spatial distributions, behaviors, and sources of PCDD/Fs in surface water and sediment from the Yangtze River Delta, Environmental Research 251 (2024) 118540.

[8] F. He, F. Wang, Y. Peng, H. Cui, G. Lv, Insight into the formation of polychlorinated dibenzo-p-dioxins and dibenzofurans in hazardous waste incineration and incinerators: Formation process and reduction strategy, Journal of Environmental Management 345 (2023) 118669.

[9] S.B. Tavakoly Sany, L. Narimani, F.K. Soltanian, R.b. Hashim, M. Rezayi, D.J. Karlen, H.N.M.E. Mahmud, An overview of detection techniques for monitoring dioxin-like compounds: latest technique trends and their applications, RSC Advances 6 (2016) 55415–55429.

[10] J. Hong, Y. Miki, K. Honda, H. Toita, Development of the automated cleanup system for the analysis of PCDDs, PCDFs and DL-PCBs, Chemosphere 88(11) (2012) 1287–1291.

[11] J.O. Lay, R. Liyanage, J.A. Gidden, THE DEVELOPMENT OF A HIGH-RESOLUTION MASS SPECTROMETRY METHOD FOR ULTRA-TRACE ANALYSIS OF CHLORINATED DIOXINS IN ENVIRONMENTAL AND BIOLOGICAL SAMPLES INCLUDING VIET NAM ERA VETERANS, Mass Spectrom Rev 40(3) (2021) 236–254.

[12] J.F. Ayala-Cabrera, M. Ábalos, E. Abad, E. Moyano, F.J. Santos, Feasibility of gas chromatography-atmospheric pressure photoionization-high-resolution mass spectrometry for the analysis of polychlorinated dibenzo-p-dioxins, dibenzofurans, and dioxin-like polychlorinated biphenyls in environmental and feed samples, Anal Bioanal Chem 412(15) (2020) 3703–3716.

[13] E.J. Reiner, The analysis of dioxins and related compounds, Mass Spectrom Rev 29(4) (2010) 526–59.

[14] G. Otarola, H. Castillo, S. Marcellini, Aryl hydrocarbon receptor-based bioassays for dioxin detection: Thinking outside the box, J Appl Toxicol 38(4) (2018) 437–449.

[15] S. Safe, U.H. Jin, H. Park, R.S. Chapkin, A. Jayaraman, Aryl Hydrocarbon Receptor (AHR) Ligands as Selective AHR Modulators (SAhRMs), Int J Mol Sci 21(18) (2020).

[16] R. Pohjanvirta, M. Viluksela, Novel Aspects of Toxicity Mechanisms of Dioxins and Related Compounds, Int J Mol Sci 21(7) (2020).

[17] L. Sládeková, S. Mani, Z. Dvořák, Ligands and agonists of the aryl hydrocarbon receptor AhR: Facts and myths, Biochemical Pharmacology 213 (2023) 115626.

[18] C.A. Opitz, P. Holfelder, M.T. Prentzell, S. Trump, The complex biology of aryl hydrocarbon receptor activation in cancer and beyond, Biochemical Pharmacology 216 (2023) 115798.

[19] A.J. Murk, J. Legler, M.S. Denison, J.P. Giesy, C. van de Guchte, A. Brouwer, Chemical-activated luciferase gene expression (CALUX): a novel in vitro bioassay for Ah receptor active compounds in sediments and pore water, Fundam Appl Toxicol 33(1) (1996) 149–60.

[20] C. Budin, H. Besselink, B.M.A. van Vugt-Lussenburg, H.Y. Man, B. van der Burg, A. Brouwer, Induction of AhR transactivation by PBDD/Fs and PCDD/Fs using a novel human-relevant, high-throughput DR(human) CALUX reporter gene assay, Chemosphere 263 (2021) 128086.

[21] C.A. Miller, A human aryl hydrocarbon receptor signaling pathway constructed in yeast displays additive responses to ligand mixtures, Toxicology and applied pharmacology 160 3 (1999) 297–303.

[22] T. Xu, A. Young, E. Marr, G. Sayler, S. Ripp, D. Close, A rapid and reagent-free bioassay for the detection of dioxin-like compounds and other aryl hydrocarbon receptor (AhR) agonists using autobioluminescent yeast, Anal Bioanal Chem 410(4) (2018) 1247–1256.

[23] C.G. England, E.B. Ehlerding, W. Cai, NanoLuc: A Small Luciferase Is Brightening Up the Field of Bioluminescence, Bioconjug Chem 27(5) (2016) 1175–1187.

[24] L. Yi, M.C. Gebhard, Q. Li, J.M. Taft, G. Georgiou, B.L. Iverson, Engineering of TEV protease variants by yeast ER sequestration screening (YESS) of combinatorial libraries, Proc Natl Acad Sci U S A 110(18) (2013) 7229–34.

[25] H. Li, L. Dong, J.P. Whitlock, Jr., Transcriptional activation function of the mouse Ah receptor nuclear translocator, J Biol Chem 269(45) (1994) 28098–105.

[26] M. Kawanishi, K. Ohnisi, H. Takigami, T. Yagi, Simple and rapid yeast reporter bioassay for dioxin screening: evaluation of the dioxin-like compounds in industrial and municipal waste incineration plants, Environ Sci Pollut Res Int 20(5) (2013) 2993–3002.

[27] P. Leskinen, K. Hilscherova, T. Sidlova, H. Kiviranta, P. Pessala, S. Salo, M. Verta, M. Virta, Detecting AhR ligands in sediments using bioluminescent reporter yeast, Biosensors and Bioelectronics 23(12) (2008) 1850–1855.

[28] T.D. Hubbard, I.A. Murray, W.H. Bisson, A.P. Sullivan, A. Sebastian, G.H. Perry, N.G. Jablonski, G.H. Perdew, Divergent Ah Receptor Ligand Selectivity during Hominin Evolution, Mol Biol Evol 33(10) (2016) 2648–58.

[29] P.A. Harper, C.L. Golas, A.B. Okey, Characterization of the Ah receptor and aryl hydrocarbon hydroxylase induction by 2,3,7,8-tetrachlorodibenzo-p-dioxin and benz(a)anthracene in the human A431 squamous cell carcinoma line, Cancer Res 48(9) (1988) 2388–95.

[30] Y. Zhao, D. Li, Z. Zhang, L. Pan, In vitro recombinant yeast assay reveals the binding of 2,3,7,8-tetrachlorodibenzo-p-dioxin (TCDD) and aryl hydrocarbon receptor (AhR) from scallop Chlamys farreri, Toxicol In Vitro 59 (2019) 64–69.

