## Supplementary material for "A new yeast-based bioreporter for simple, sensitive, and cost-effective detection of dioxin-like compounds": Table S1;Figure S1.

Author Contributions:

\# Yingying Liang, Hailin Liua and Lin Wang contributed equally

*Corresponding author at:

Sijing Jiang, School of Life Sciences, Hubei University, Wuhan, Hubei 430062 China

Zhenghui Lu, School of Life Sciences, Hubei University, Wuhan, Hubei 430062 China

Guimin Zhang, College of Life Science and Technology, Beijing University of Chemical Technology, Beijing 100029, China

**Supplementary Table**

**Table S1. The main primers used in the experiment.**

| Primer | Sequence |
| --- | --- |
| ARNT -F | GTTAATTAACTGCAGGAATTCATCATTCTGAAAAGGGGGGAAACATAGTTAG |
| ARNT -R | GCAGTCAACCCTCACTAAAGGGCGGCCGCACATGGCGGCGACTACTGCC |
| AhR -F | CTATACTTTAACGTCAAGGAGAAAAAACATGAACAGCAGCAGCGCC |
| AhR -R | GTTATCAGATCTCGAGGGATCCGGGTTACAGGAATCCACTGGATGTCAAATC |
| XRE5 -F | GAAGAATTGTTAATTAACTGCAGTTGGATCCTTGCGTGACAATTC |
| TCYC1-R | CTAAAGGGCGGCCGCACTGAATTCGCAAATTAAAGCCTTCGAGCG |
| G418-F | GATTTGACATCCAGTGGATTCCTGTAATCATGTAATTAGTTATGTCACGC |
| G418-R | GTGTTGGTTTTTATATGTTTTCAGTATAGCGACCAGCATTCAC |
| H-up-F | GTTAATTAACTGCAGGAATTCCTCGAGGGATTGAGGCCACAGCAAGACCGGCC |
| H-up-R | CTATGTTTCCCCCCTTTTCAGAATAAGTTATGTATTGTTTATTTTCCCTTTAATTTTAG |
| H-down-F | GTGAATGCTGGTCGCTATACTGAAAACATATAAAAACCAACACAATAAAAAAAAGG |
| H-down-R | CAGATCTCGAGGGATCCCTCGAGTCTTAGTTGGTAGCACTTTGATGAG |
| YZ-R | GATGGACTGGCCAGCTACAATC |
| YZ-F | GGTAGCGTTGCCAATGATGTTAC |
| Nluc-F | GATGGACTGGCCAGCTACAATC |
| Nluc-R | CATGATTACGCCAGAATGCGTTC |
| ARNT-1-F | GTCAAGGAGAAAAAACCCCGGATCCCTCGAATGGCGGCGACTACTGCC |
| ARNT-1-R | CTCACTATAGGGCGAATTGGTTATTCTGAAAAGGGGGGAAACATAGTTAG |
| AhR381-F | GGAAGACCAGATTATATCATTGCCACTCAGAGACCACTAACAG |
| AhR381-R | ATCTGTTAGTGGTCTCTGAGTGGCAATGATATAATCTGGTCTTCC |

**Supplementary Figure**

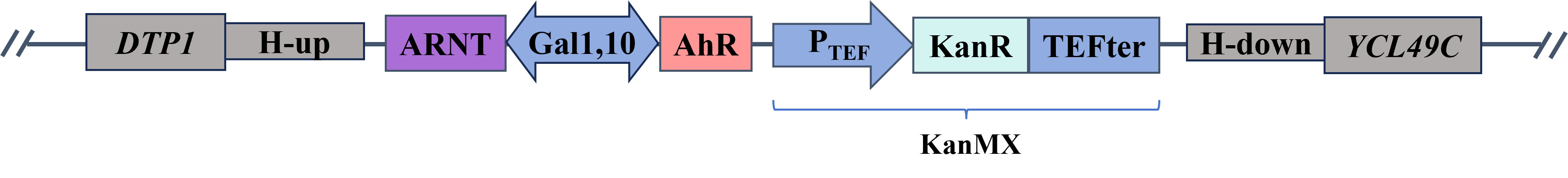

**Figure S1. Sensor module integration diagram**
